# Lack of support for Deuterostomia prompts reinterpretation of the first Bilateria

**DOI:** 10.1101/2020.07.01.182915

**Authors:** Paschalia Kapli, Paschalis Natsidis, Daniel J. Leite, Maximilian Fursman, Nadia Jeffrie, Imran A. Rahman, Hervé Philippe, Richard R. Copley, Maximilian J. Telford

## Abstract

The bilaterally symmetric animals (Bilateria) are considered to comprise two monophyletic groups, Protostomia and Deuterostomia. Protostomia contains the Ecdysozoa and the Lophotrochozoa; Deuterostomia contains the Chordata and the Xenambulacraria (Hemichordata, Echinodermata and Xenacoelomorpha). Their names refer to a supposed distinct origin of the mouth (stoma) in the two clades, but these groups have been differentiated by other embryological characters including embryonic cleavage patterns and different ways of forming their mesoderm and coeloms. Deuterostome monophyly is not consistently supported by recent studies. Here we compare support for Protostomia and Deuterostomia using five recently published, phylogenomic datasets. Protostomia is always strongly supported, especially by longer and higher quality genes. Support for Deuterostomia is always equivocal and barely higher than support for paraphyletic alternatives. Conditions that can cause tree reconstruction errors - inadequate models, short internal branch, faster evolving genes, and unequal branch lengths - correlate with statistical support for monophyletic deuterostomes. Simulation experiments show that support for Deuterostomia could be explained by systematic error. A survey of molecular characters supposedly diagnostic of deuterostomes shows many are not valid synapomorphies. The branch between bilaterian and deuterostome common ancestors, if real, is very short. This finding fits with growing evidence suggesting the common ancestor of all Bilateria had many deuterostome characteristics. This finding has important implications for our understanding of early animal evolution and for the interpretation of some enigmatic Cambrian fossils such as vetulicolians and banffiids.

## Main

In the case of a burst of diversification such as is apparent at the base of the Cambrian, reconstructing the evolutionary history of taxa is difficult. Limited time between speciations (short internal branches) lead to the accumulation of a small amount of phylogenetic signal and this signal is easily blurred by the numerous substitutions occurring later (long terminal branches). These problems are exacerbated by heterogeneities in the evolutionary process which can result in model violations that can cause systematic errors^1^. Very short internal branches also strongly imply minimal evolutionary change meaning that common ancestors of nested clades are expected to have been highly similar.

### The deuterostome branch has weak support

The different topologies relating Chordata, Xenambulacraria and Protostomia seen in recently published animal phylogenies suggest that the support for monophyletic deuterostomes (Fig. 1a) may be low, in contrast with the consistent, strong support for the monophyly of the protostomes. To study this difference, we have gathered five recent, independently generated, phylogenomic datasets covering the diversity of animal phyla (i.e., ^2–6^). We use these data to investigate the support for different topologies relating the Chordata, Xenambulacraria, Ecdysozoa and Lophotrochozoa. We first asked whether the branches leading to protostomes and deuterostomes were of similar length, under the assumption that both are monophyletic.

**Figure 1.**
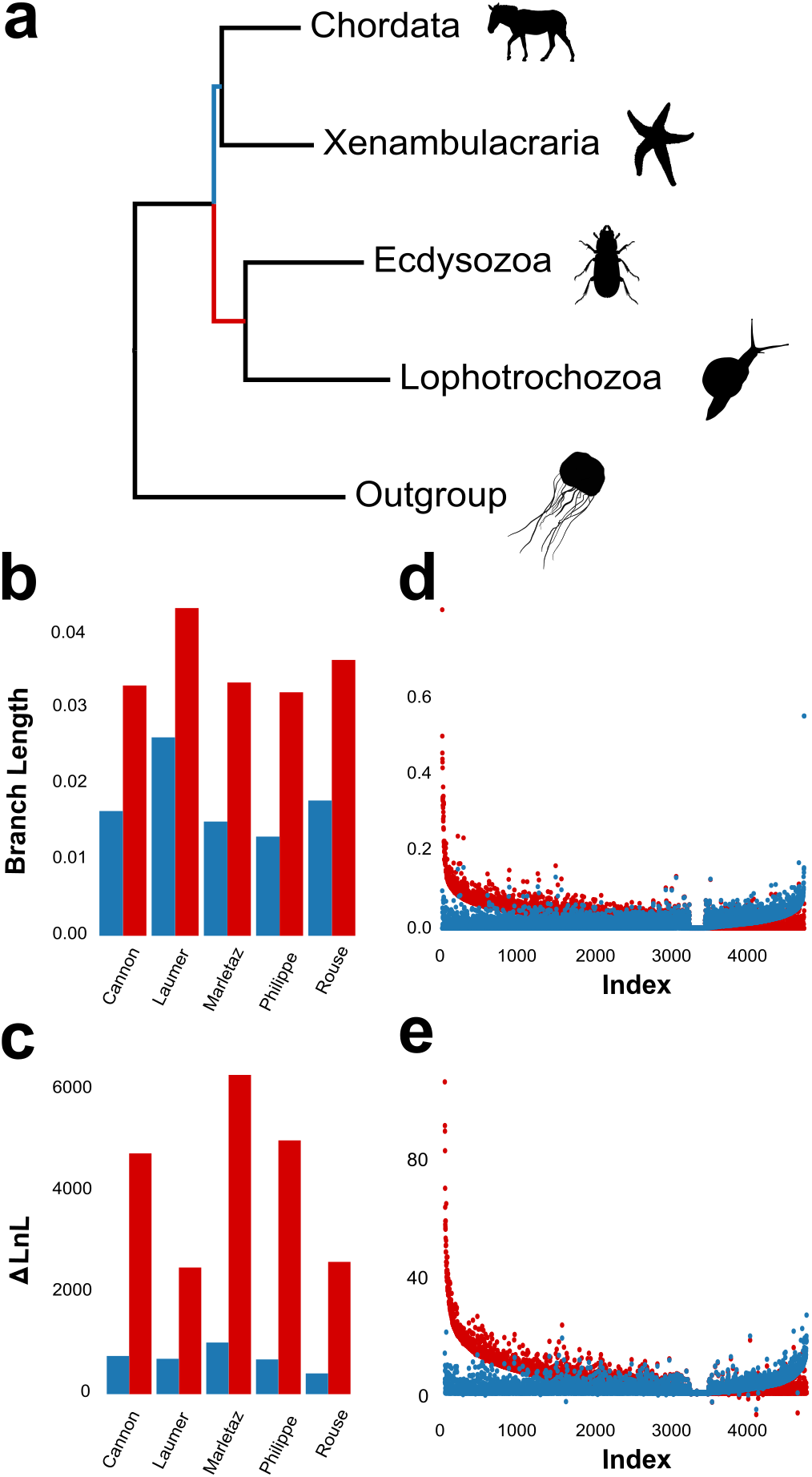
The deuterostome branch is short and weakly supported compared to the protostome branch. a. Canonical tree showing relationships between major metazoan clades. The Bilateria consists of two branches, Protostomia (Red: Lophotrochozoa and Ecdysozoa) and Deuterostomia (Blue: Chordata and Xenambulacraria). b. Comparison of branch lengths from the bilaterian common ancestor to the deuterostome common ancestor (blue) and to the protostome common ancestor (red). In five independently curated large datasets the protostome branch is twice as long as the deuterostome branch. c. Comparison of estimated protostome and deuterostome branch lengths for individual genes. Genes from all five datasets are ordered by the difference in length between protostome and deuterostome branches. In 3376 out of 4826 genes (69.9%) the protostome branch is longer. In 1402 out of 4826 genes (29%) the deuterostome branch is longer. c. Comparison of decrease in lnLikelihood of a tree in which either the protostome or deuterostome branch has been collapsed (ΔlnL). In five independently curated large datasets collapsing the protostome branch reduces lnL considerably more than collapsing the deuterostome branch. cd. Comparison of estimated protostome and deuterostome branch lengths for individual genes. Genes from all five datasets are ordered by the difference in length between protostome and deuterostome branches. In 3376 out of 4826 genes (69.9%) the protostome branch is longer. In 1402 out of 4826 genes (29%) the deuterostome branch is longer. e. Comparison of protostome and deuterostome ΔlnL for individual genes. Genes from all five datasets are ordered by the difference in ΔlnL between protostomes and deuterostomes. In 3312 out of 4826 genes (68.6%) the protostome ΔlnL is larger. In 1345 out of 4826 genes (27.6%) the deuterostome ΔlnL is larger. All analyses used the LG+F+G model.

Using a fixed topology taken from the original publications for each dataset (modified when necessary to enforce monophyly of deuterostomes), we optimised branch lengths using the site-homogeneous LG+F+G model. In Figure 1b we see that, while the overall amount of change in branches leading from the bilaterian common ancestor to both protostomes and deuterostomes differs between datasets, in all cases, the branch leading to the protostomes is approximately twice the length of the branch leading to the deuterostomes. This is a conservative measure of the difference, as better fitting models (see later) give longer estimates for the protostome branch and shorter estimates for the deuterostome branch (for the Laumer dataset the protostome branch is estimated as 1.75 times longer than deuterostome under LG and 5.7 times longer under CAT-LG). This difference between protostome and deuterostome branches is reinforced when we look at the change in log likelihood (ΔlnL) between a fully resolved tree compared to trees in which either the protostome or deuterostome branch is collapsed into a polytomy. The ΔlnL observed on collapsing the branch leading to protostomes is considerably greater for all five datasets than the ΔlnL when the deuterostome branch is collapsed (Fig. 1c), showing considerably less signal supporting deuterostomes than protostomes.

To get a first indication of whether this weaker support for deuterostomes results from conflict between genes or from a lack of signal across genes and across datasets, we repeated these measures of branch lengths and ΔlnL on individual genes in all five datasets (Fig. 1d,e). We find that, for the great majority of genes across all datasets, as for the concatenated datasets, there is a longer branch leading to the protostomes than to deuterostomes (P longer: 3376 (70%); D longer: 1402 (29%), Supplementary Data 1) and consistently stronger support (larger ΔlnL) for protostomes than for deuterostomes (ΔlnL P larger: 3312 (68.6%); ΔlnL D larger: 1345 (27.8%), Supplementary Data 2). Most genes support monophyletic deuterostomes much less strongly than they support protostomes.

### Minority of genes support deuterostome monophyly

While the difference in branch lengths and likelihood indicate weaker evidence for the monophyly of deuterostomes across different genes relative to the support for protostomes, the measures presented are based on trees in which deuterostomes were constrained to be monophyletic, meaning that we could not show any topological conflict between genes. To measure conflict between genes we considered each gene individually and compared the lnLikelihood of a tree supporting protostome monophyly (“PM”) or deuterostome monophyly (“DM”) with the lnLikelihoods of two alternative topologies for each of these two clades: Protostome Paraphyly “P1”: Lophotrochozoa sister to Deuterostomia; Protostome Paraphyly “P2”: Ecdysozoa sister to Deuterostomia; Deuterostome Paraphyly “D1”: Xenambulacraria sister to Protostomia; Deuterostome Paraphyly “D2”: Chordata sister to Protostomia. (see Fig. 2). We visualised the relative lnLikelihoods for the three topologies for every gene on a triangular plot (Fig. 2). For the protostome trees, most genes have a strong preference for a single topology (77.31% on average across datasets with the LG+F+G model, Extended Data Table 1); amongst the genes that strongly prefer one topology, we see a clear majority supporting monophyletic protostomes (58.94% vs 9.71% and 8.66% for paraphyletic protostomes P1 and P2 respectively, Extended Data Table 1).

**Figure 2.**
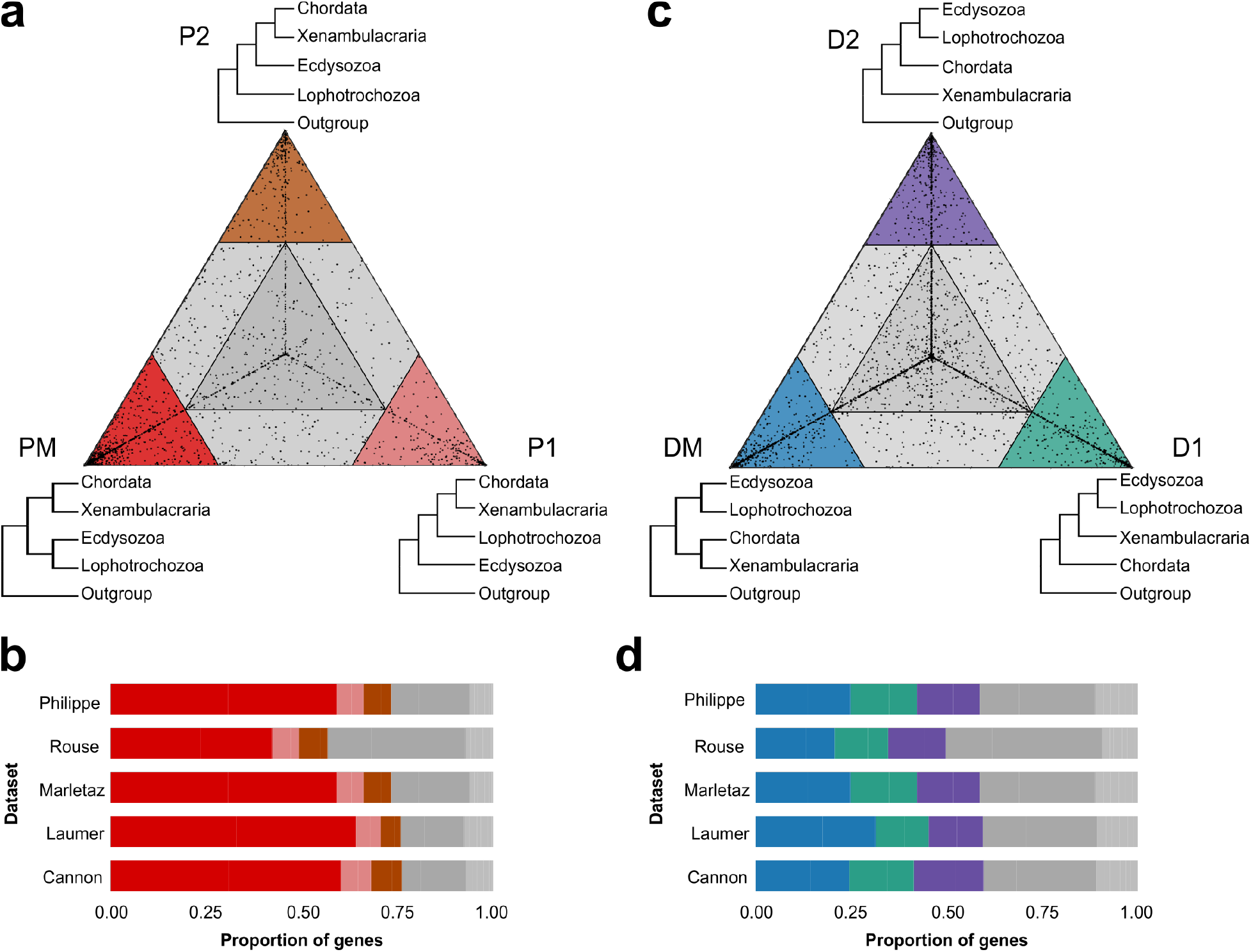
Most genes prefer monophyletic Protostomia over alternatives, few genes prefer monophyletic Deuterostomia. Triangular plots showing relative support for the three alternative topologies shown at the corners of the triangles. Each dot represents a gene and its position is determined by the relative lnLikelihoods (measured using the LG+F+G model) of the three constrained trees calculated using that gene alignment. Genes in coloured corners show a high preference for the corresponding topology. The numbers of genes found in the different coloured sectors of the large triangle after estimating likelihoods with the LG+F+G model are shown below. In the triangle plots, all genes from the 5 datasets are visualised together, in the bar plots each named dataset is represented by a bar. a. Triangle plot comparing support for monophyletic Protostomia (PM) versus two alternative topologies with paraphyletic Protostomia (P1, P2). b. Bar plot showing that across five datasets the majority of genes strongly prefer the monophyletic Protostomia topology. c. Triangle plot comparing support for monophyletic Deuterostomia (DM) versus two alternative topologies with paraphyletic Deuterostomia (D1, D2). d. Bar plot showing that across five datasets a minority of genes strongly prefer the monophyletic Deuterostomia topology over paraphyletic topologies or the grey areas (representing weak or no preference).

For the deuterostomes, the results are more equivocal; fewer genes have strong preference for a single topology (66.99% on average, Extended Data Table 2) and, amongst genes that do prefer one topology, there is only a small excess in the number of genes strongly supporting monophyletic deuterostomes over the other two topologies (28.65% vs 19.17% and 19.17% for paraphyletic deuterostomes D1 and D2 respectively, Extended Data Table 2). Overall, these experiments show that, while the protostome clade is strongly supported over either of the two topologies with paraphyletic protostomes, the same is not true for monophyletic deuterostomes which have a very small advantage over either of the alternative topologies.

For all five datasets, the gene alignments strongly supporting the monophyletic protostomes topology were longer and had higher monophyly scores (a measure of their ability to support known clades) compared to the gene alignments that supported either of the two alternative topologies (Extended Data Fig. 1). For the deuterostomes, there was minimal evidence of a significant difference of alignment length between genes supporting DM versus D1 or D2 (Extended Data Fig. 1) and there were no significant differences in monophyly scores between genes supporting DM versus D1 or D2 (Extended Data Table 3). Monophyletic Protostomia, but not monophyletic Deuterostomia, is strongly preferred across five independent datasets by a clear majority of individual genes which are on average longer and have a stronger phylogenetic signal.

### Deuterostome monophyly could result from systematic error

Whatever the true topology relating Xenambulacraria, Chordata and Protostomia, we have shown that the branch separating them is short and might therefore be especially sensitive to systematic errors. Systematic errors are generated by heterogeneities in the process of substitution that are not accommodated by the models used and are exacerbated in faster evolving datasets. In the context of unequal rates of evolution amongst taxa this may lead to a long branch attraction (LBA) artefact^7^. We next consider the possibility that monophyletic Deuterostomia could result from systematic error.

For each dataset we measured the average branch lengths within each of the protostome, chordate and xenambulacrarian clades (average distance from bilaterian common ancestor to each taxon within each clade) and the average distance from each outgroup taxon to the bilaterian common ancestor. The branches leading to the outgroups and to the protostomes are always longer than the branch leading to the deuterostomes (Fig. 3a). Under LBA, the long-branched protostomes and outgroup might be expected to be mutually attracted resulting in clustering of short branched Chordata and Xenambulacraria. We note that the three datasets which supported monophyletic deuterostomes^2,3,6^ had the largest rate inequalities with the longest average protostome and outgroup branch lengths relative to chordate and xenambulacrarian branches (Fig. 3a).

**Figure 3.**
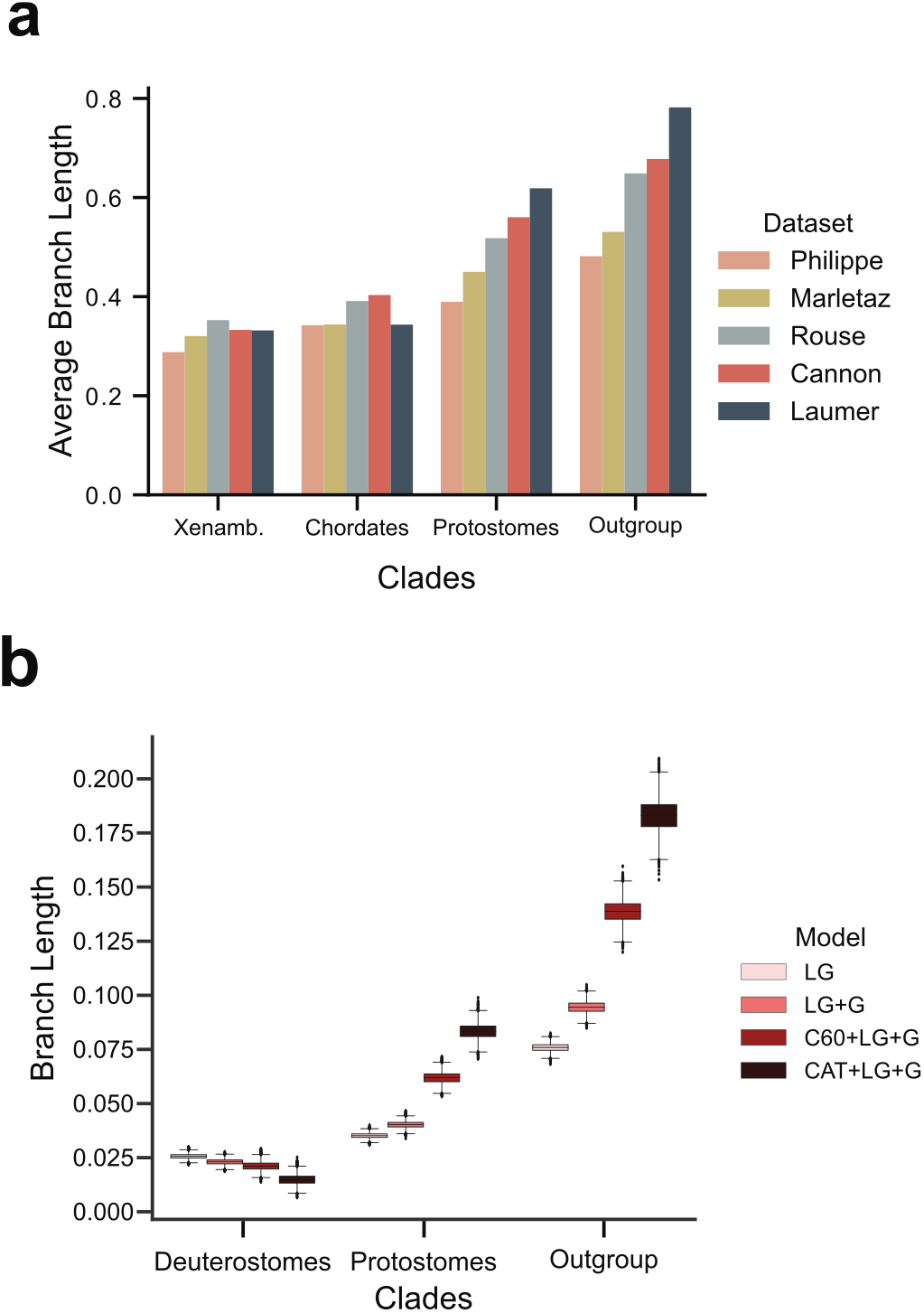
Evidence from empirical data for systematic error supporting monophyletic Deuterostomia. a. Bar chart showing the average branch lengths within Chordata, Xenambulacraria, Protostomia and from outgroup taxa to the root of the Bilateria. Datasets with the greatest branch length heterogeneity (Rouse, Cannon, Laumer) are those that support deuterotome monophyly. b. Box plot showing the lengths of the branches leading to the Deuterostomia, Protostomia and from outgroups to Bilateria estimated using different models of evolution and 50,000 sites, 36 species from the Laumer dataset. Cross validation shows that models fit the data increasingly well in the order LG < LG+G < C60+LG+G < CAT+LG+G. With improving model fit, the estimated length of the deuterostome branch decreases, and estimated lengths of the protostome and outgroup branches increase. Model violations (e.g. using site homogeneous models such as LG) can lead to an underestimate of the likelihood of change on long branches. This would be predicted to promote LBA between the protostomes and outgroup and to give artifactual support to deuterostomes. Less well-fitting models tend to support monophyletic deuterostomes.

To compare the relative rates of the 5 datasets we removed the effect of taxon sampling by reducing all 5 datasets to the 8 species all have in common and measured the rate of substitution across the tree; the three datasets that support monophyletic Deuterostomia are the fastest evolving (substitutions per site across the 8 taxon tree - Philippe: 1.971, Marletaz: 2.2082, Cannon: 2.537, Laumer: 2.6578, Rouse: 2.8741).

Finally, while two datasets support paraphyletic deuterostomes under site heterogeneous models^4,5^ which account for across site amino acid preference variability^8^, all five datasets support monophyletic deuterostomes when using site homogeneous models. Unequal rates, faster evolving loci and inadequate site homogeneous models are all known to promote LBA artifacts, especially in the context of short internal nodes^1,7,9^ and all these conditions correlate with support for Deuterostomia.

We further examined the possible effect of model misspecification by comparing branch lengths estimated using different models. We used the Laumer et al. (2019)^3^ dataset which has the clearest branch length heterogeneity (Fig. 3a). For computational speed, we selected 36 taxa focussing on maintaining the taxonomic diversity of the full dataset and excluding taxa with a large percentage of missing data. We randomly selected 50,000 sites from the full alignment of 106,186 sites. We conducted cross validation comparing 4 increasingly complex models (LG, LG+G, C60+LG+G and CAT+LG+G). As expected less complex models have significantly worse fit to the data than the more complex (average cross validation scores LG: −74877.78, LG+G: −70348.95, C60+LG+G: −68707.89, CAT+LG+G: −68031.36). We estimated the lengths of the branches leading to the deuterostomes, the protostomes and the outgroup with these different models. Using models that fit the data increasingly well, we see that the estimated length of the deuterostome branch decreases, and the length of the protostome and outgroup branches increase (Fig. 3b). Using inadequate models could therefore underestimate the likelihood of convergence between Protostomia and outgroup. This would be predicted to result in a long branch attraction between protostomes and outgroups resulting in monophyletic deuterostomes.

To evaluate further the idea that model misspecification in the context of unequal evolutionary rates between taxa might lead to a long branch artifact, we followed a data simulation approach. We ask whether any of the three topologies relating the Protostomia, Chordata and Xenambulacraria (DM, and the two paraphyletic alternatives D1 and D2, Fig. 4a) could gain artifactual support as a result of systematic error. Using the 50,000 positions and 36 taxa from the Laumer dataset, we estimated parameters using the three alternative topologies (DM, D1, D2) under the best fitting CAT+LG+G model implemented in phylobayes^10^ (Fig. 4a). For each topology we simulated 100 datasets using the parameters we had estimated.

**Figure 4.**
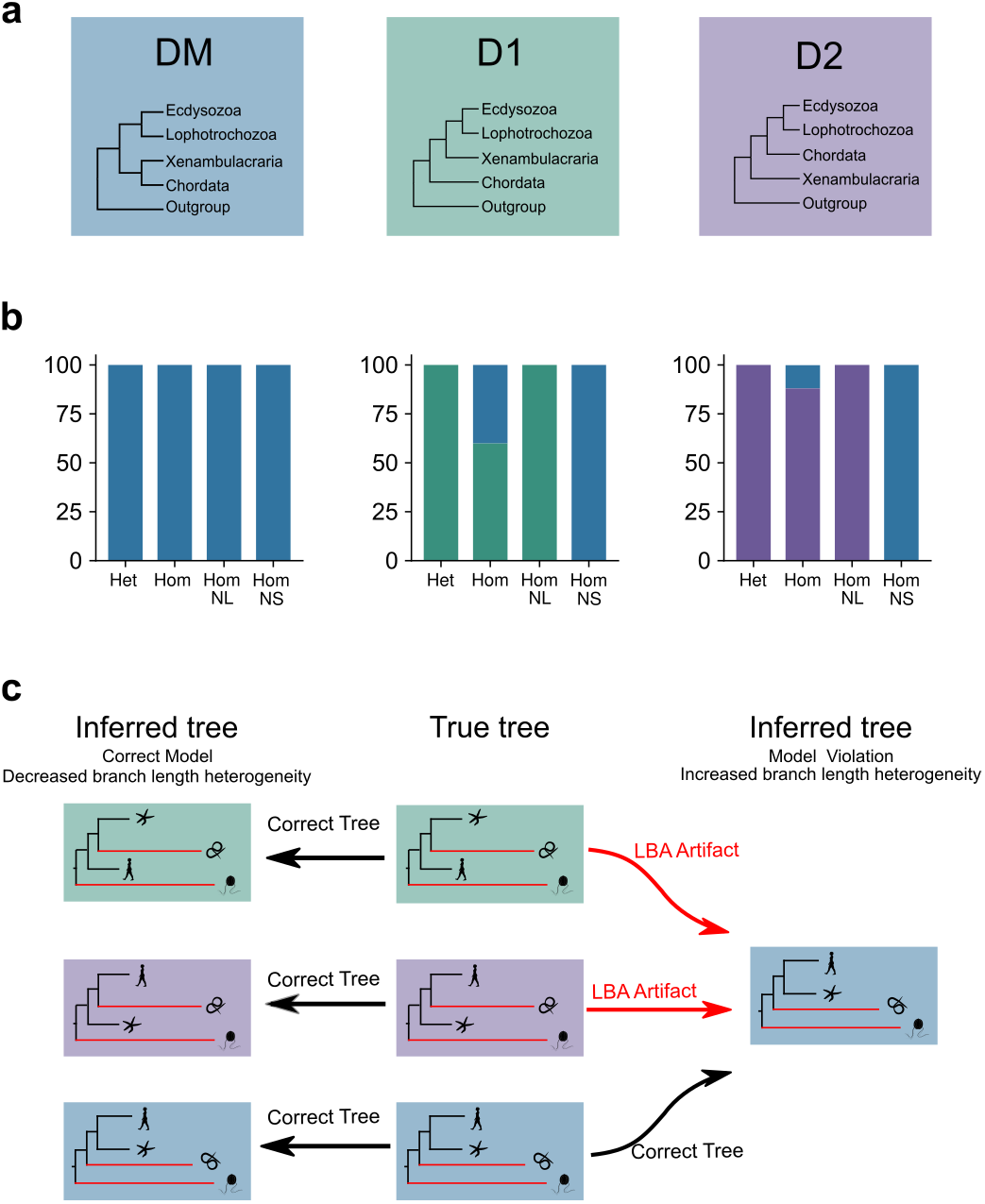
Simulations show model violations and unequal rates result in artifactual support for monophyletic Deuterostomia. a. 100 datasets were simulated using parameters estimated under a site heterogeneous model (CAT+LG+G) for each of the three topologies shown (coloured boxes). b. For each simulated dataset a maximum likelihood tree was reconstructed under four conditions: Site heterogeneous C60+LG+F+G model (Het); model violating site homogenous LG+F+G model (Hom); homogeneous model with long branch protostomes and outgroup taxa removed (HomNL); homogeneous model with short branch protostomes and outgroup taxa removed (HomNS). For each condition, the number of times the three possible topologies were reconstructed is shown in the bar charts - the colour indicates the topology supported. Data simulated under the DM topology always yield the correct topology under all conditions. Under the site heterogeneous model, D1 and D2 data always yield a correct topology. Under the model violating site homogeneous model, D1 and D2 data yield an incorrect topology in 40% and 12% of replicates respectively. The incorrect tree is always DM. Under the site homogeneous model with long branch protostomes and outgroups removed (reducing LBA) D1 and D2 data always yield a correct topology. Under the site homogeneous model with short branch protostomes and outgroups removed (enhancing LBA) D1 and D2 data always yield an incorrect topology. The incorrect tree is always DM. c. Interpretation of simulation results as an LBA artifact. The tree topologies under which data were simulated (true tree) are shown in the middle. The long branches leading to the protostomes and outgroups are indicated in red. Under conditions that minimise systematic error (Left: site heterogeneous model or reduction of branch length heterogeneity by removing long branches) the correct tree (DM, D1 or D2) is always reconstructed. Under conditions that enhance systematic error (Right: model violating site homogeneous models or increase of branch length heterogeneity by removing short branches) the DM topology is reconstructed using data simulated under all three topologies.

For data simulated according to the DM topology (Chordata plus Xenambulacraria) we are able to reconstruct monophyletic deuterostomes (DM) correctly in 100% of replicates under both correct (site heterogeneous) and model violating (site homogeneous models) (Fig. 4b). In contrast, for the data simulated under the two alternative hypotheses (D1, D2), recovering the true relationships was more challenging. As predicted by the consistency of probabilistic models, under the correct site heterogeneous model the correct topology was always recovered. Under the model-violating site homogeneous model, which would be expected to exacerbate LBA artifacts, the true tree was recovered in 60% and 88% of the datasets simulated under the D1 and D2 hypotheses respectively (Fig. 4b). In both cases, the remaining 40% (D1) and 12% (D2) of the datasets incorrectly recovered monophyletic deuterostomes (the DM topology). This result suggests that if either D1 or D2 were the true tree, LBA stemming from a combination of the long branches leading to protostomes and outgroups, short internal branches and model misspecification, could result in artifactual support for deuterostome monophyly.

To explore the hypothesis of unequal rates of evolution lending exaggerated support to the monophyletic Deuterostomia topology (DM), we re-inferred trees using the same simulated data after first removing the 13 longest branched taxa of the protostome and outgroup clades, a commonly adopted strategy to reduce LBA artifacts^11,12^. We re-inferred the trees using the poorly fitting site homogeneous model (LG+F+G) which had resulted in some incorrect topologies with the full dataset. Regardless of the topology under which the data were simulated (DM, D1 or D2) removing these long branches resulted in recovering the correct topology in 100% of cases (Fig. 4b). Despite the model violation there was no long branch attraction artifact.

We contrast this result with an equivalent experiment designed to exaggerate any artifact caused by rate heterogeneity. We removed the 13 shortest protostome and outgroup taxa from the simulated data and again re-inferred the tree topologies under the site homogeneous LG+F+G model. Regardless of the topology under which the data were simulated (DM, D1 or D2), we recovered monophyly of the deuterostomes (DM topology) in 100% of replicates (Fig. 4b). These results show that, if deuterostomes are paraphyletic (either D1 or D2 topology), then conditions we have shown to affect real datasets could easily result in artifactual support for monophyletic deuterostomes (Fig. 4c).

### Reappraising deuterostome molecular synapomorphies

Studies of different classes of molecular characters also suggest stronger support for Prostostomia than Deuterostomia. Protostomes have 12 unique microRNA families^13^ compared to 1 in deuterostomes and protostomes share 58 ‘near intron pairs’ compared to just 7 in deuterostomes. Protostomes also share a highly distinct, conserved variant of the mitochondrial NAD5 protein ^14^ and a hidden break in their 28S ribosomal RNA^15^.

In their comparison of hemichordate and chordate genomes, Simakov et al.^16^ conducted a systematic search for genes unique to the deuterostomes. Using the greater number and diversity of sequenced genomes now available, we have identified orthologs of many of these genes in non-bilaterian metazoans and/or in protostomes. Overall, we have found evidence for 20 out of 31 of these ‘deuterostome novelties’ in protostomes and/or non bilaterian metazoans suggesting they are bilaterian plesiomorphies (Supplementary Data 4). Equivalent searches for characters unique to either Chordata plus Protostomia or Xenambulacraria plus Protostomia have not been conducted and it is therefore not clear how to interpret the 11 remaining Chordata plus Xenambulacraria specific characters. If deuterostomes are not monophyletic, these 11 genes must have been lost in protostomes.

Simakov et al. also describe a deuterostomian ‘pharyngeal gene cluster’^16^. This is a micro-syntenic block of 7 genes (nkx2.1, nkx2.2, msxlx, pax1/9, slc25A21, mipol1 and foxA), found complete only in Ambulacraria and Chordata. Several of these genes have functional links to pharynx patterning and the formation of pharyngeal slits. Different protostomes do have some of these genes linked (nkx2.1 nkx2.2 - various protostomes); nkx2.2 msxlx (*Lottia*); pax1/9 slc25A21 (*Lottia*); mipol1 foxA (various protostomes) and pax1/9 foxA (distant but on the same chromosome in *Caenorhabditis*)^17^. Msxlx is adjacent to 3 nkx2.1/2-type genes in *Trichoplax* and is also relatively close to foxA in the genomes of medusozoan and octocoral cnidarians (cnidarians lack pax1/9; msxlx is also absent in hexacorals). We have also identified a linkage between msxlx and pax1/9 in the genome of the protostome *Phoronis australis* (separated by 0.34 mb, with 17 intervening genes) (Supplementary Data 4). These links raise the possibility that at least five (nkx2.1 nkx2.2 msxlx pax1/9 slc25A21) and perhaps all of the genes in the cluster were linked in the bilaterian common ancestor and that the cluster is a bilaterian character that has been dispersed in different protostome lineages (Fig. 5a).

**Figure 5.**
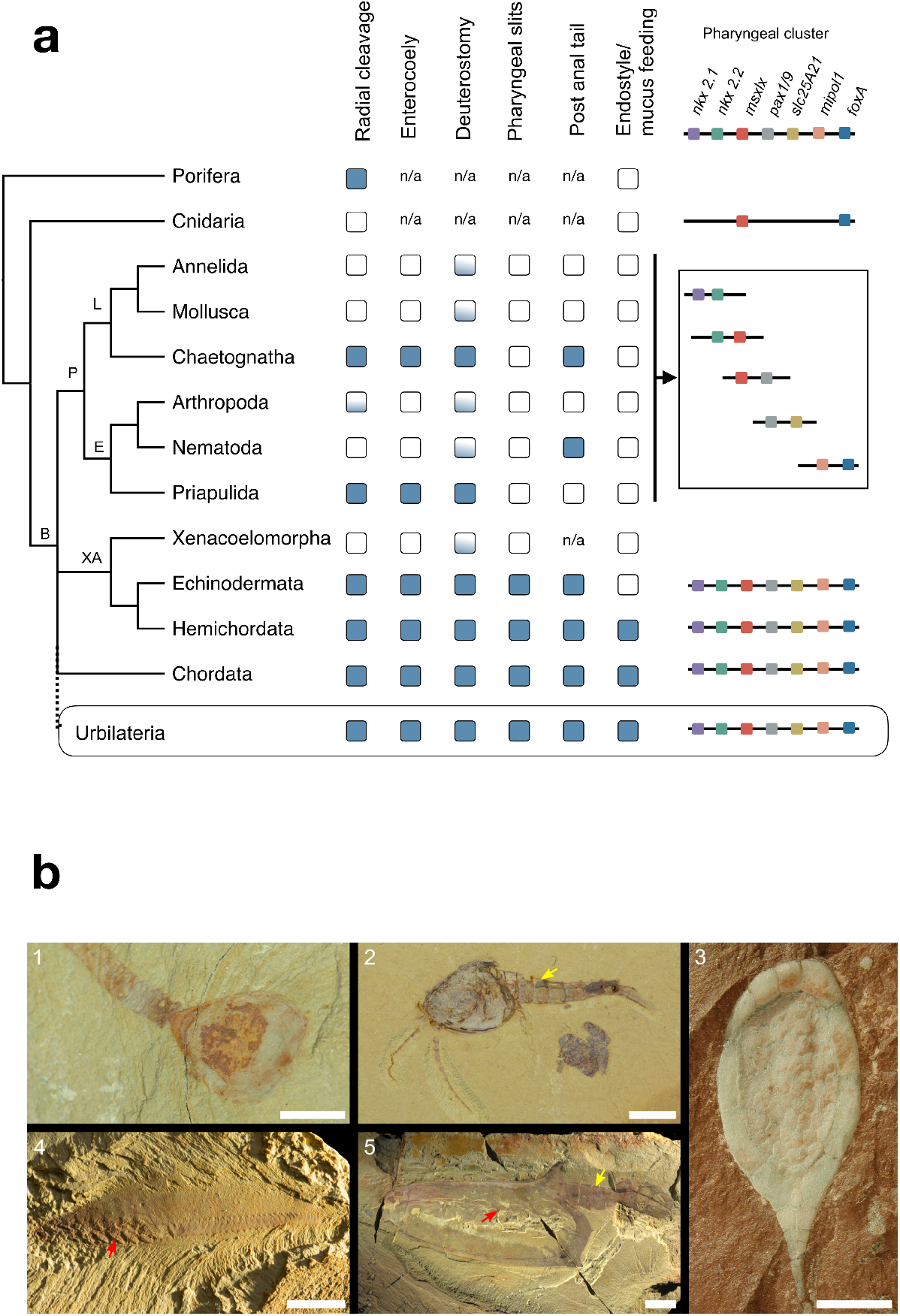
Character evolution implied by a short or non-existent deuterostome branch. a. Distribution of characters on a phylogenetic tree with an unresolved polytomy at the base of the Bilateria involving Protostomia (P), Xenambulacraria (XA) and Chordata. (L = Lophotrochozoa; E = Ecdysozoa; B = Bilateria). Presence or absence of the characters discussed in the main text are indicated (blue box = present; white box = absent; blue/white box = variable; n/a = not applicable). The pharyngeal cluster contains the seven genes indicated by coloured boxes. The cluster has not been found intact in any single protostome, but most possible pairs/triplets of adjacent genes are found linked in one or more protostomes or non-bilaterian metazoans (box represents various protostomes see text for details), implying the common ancestor of protostomes had an intact cluster. Urbilateria is the common ancestor of all three clades (dotted line) and its characteristics can be inferred as those present in Chordata and Xenambulacraria (and, for some characters, in Protostomia). All characters previously considered to be diagnostic of deuterostomes are likely to have been present in Urbilateria as indicated. b. Cambrian bilaterians. 1, The lophotrochozoan *Lingulella chengjiangensis* (Cambrian Series 2, Yunnan Province, China). 2, The ecdysozoan *Chuandianella ovata* (Cambrian Series 2, Yunnan Province, China). 3, The xenambulacrarian *Protocinctus mansillaensis* (Cambrian Series 3, Spain). 4, The chordate *Myllokunmingia fengjiaoa* (Cambrian Series 2, Yunnan Province, China). 5, The problematic bilaterian *Vetulicola cuneata* (Cambrian Series 2, Yunnan Province, China). Pharyngeal slits (red arrows) are present in *Vetulicola* and *Myllokunmingia*, while a segmented bipartite body (yellow arrows) is a feature of *Vetulicola* and *Chuandianella*. If Urbilateria possessed pharyngeal slits, as discussed in the main text, *Vetulicola* could represent a stem protostome. Images 1, 2, 4 and 5 courtesy of Yunnan Key Laboratory for Palaeobiology and MEC International Joint Laboratory for Palaeobiology and Palaeoenvironment, Yunnan University, Kunming, China. Image 3 courtesy of Samuel Zamora. Scale bars: 5 mm (1–3); 10 mm (4, 5).

## Discussion

We have shown that the branch leading to monophyletic Deuterostomia, if it exists, is short and weakly supported; for comparison we demonstrate unequivocal strong support for Protostomia especially from higher quality gene alignments. Comparisons of rates and relative branch lengths between data sets and the effects of model misspecification show that support for monophyletic Deuterostomia correlates with conditions expected to enhance systematic error. The contention that support for monophyletic Deuterostomia could result from systematic error is supported by our simulation experiments. Systematic error is especially likely to affect our ability to reconstruct relationships between taxa separated by the very short branches we have described.

The short branches and gene-tree heterogeneity we observe suggest that Chordata, Xenambulacraria and Protostomia emerged from two closely spaced speciation events. The gene-tree discordance could reflect true differences in the evolutionary histories of genes resulting from incomplete lineage sorting^18^. Testing this will require the development of methods that accommodate the LBA-causing heterogeneities affecting these data^19^.

Such a short deuterostome branch has important consequences for our understanding of character evolution at the base of the Bilateria as (assuming it exists) it implies a short period of time separating the last bilaterian common ancestor (Urbilateria) and the last deuterostome common ancestor (Urdeuterostomia)^4,20^. Hybridisation between lineages following speciation is another process that would blur the distinction between these ancestral taxa. If the deuterostomes are paraphyletic, as some of our analyses suggest, then the last common ancestor of Chordata and Xenambulacraria was the last common ancestor of the Bilateria meaning Urbilateria was a deuterostome and characters common to Xenambulacraria and Chordata are bilaterian plesiomorphies.

The idea that Urbilateria possessed some deuterostome characters is not new. Grobben included the phylum Chaetognatha in his Deuterostomia (Grobben 1908)^21^. Chaetognaths are now known to be lophotrochozoan protostomes^5,22^ but have the three main deuterostome characters^23^ of radial cleavage, forming their coeloms and mesoderm by enterocoely and their anus forms in the vicinity of the closed blastopore - the mouth is a secondary opening - and by this definition they are deuterostomes. Deuterostomy and radial cleavage are also found in the Ecdysozoa as observed in priapulid worms^24,25^. Radial cleavage, enterocoely and deuterostomy are characters found in both protostomes and deuterostomes and are therefore parsimoniously reconstructed as primitive characteristics of the Bilateria.

Members of both Chordata and Xenambulacraria also have pharyngeal slits, a post anal tail and a form of endostyle (a pharyngeal tissue that secretes iodine-rich mucus for filter feeding)^26^. Our results suggest these were also characteristics of Urbilateria (rather than deuterostome synapomorphies) implying they have been altered beyond recognition or lost in the lineage leading to the protostomes. Our evidence that the pharyngeal cluster of genes likely existed in the protostome ancestor implies that stem protostomes may have had pharyngeal slits.

If Urbliateria had these deuterostome characters there are major implications for understanding early animal evolution. A number of fossil forms from the Cambrian Period have been interpreted as stem deuterostomes because, while lacking most defining characteristics of extant deuterostome phyla, they appear to possess pharyngeal slits^27–32^ (Fig. 5b). The presence of pharyngeal slits in vetulicolians^27,29,30^, has been used as evidence for placing them as stem- or total-group deuterostomes, with the bipartite vetulicolian body plan taken as a model for the ancestral deuterostome^27,30,33^. If pharyngeal slits were present in Urbilateria, this removes the key character supporting placement of vetulicolians with deuterostomes and the presence in vetulicolians of a terminal anus and segmentation^30^ could indicate that they are stem protostomes that had lost a post anal tail but retained pharyngeal slits. Banffiids, which lack pharyngeal slits but are otherwise morphologically similar to vetulicolians^32^ (Conway Morris et al., 2015), might occupy a more derived position in the protostome stem group.

This suite of complex characters in Urbilateria also suggests a relatively large, filter-feeding animal arguing against a small and simple urbilaterian^2,34^. A large Urbilateria would suggest that the apparent lack of Precambrian bilaterians cannot be easily explained as the result of poor preservation potential due to small size and simple morphology, and emphasises the gap between molecular clock estimates for the origin of bilaterians and their oldest fossil evidence^35^.

While we have shown that support for monophyletic Deuterostomia is exaggerated by systematic errors, we have not attempted definitively to resolve this polytomy. The conclusions we have reached as regards character evolution are, however, unlikely to be affected by resolving the relationships between these extremely short branches. Nevertheless, recent analyses employing large datasets and the best available models give tentative support to a sister group relationship between the Chordata and Protostomia to the exclusion of the Xenambulacraria. Based on the shared character of a centralised nervous system in Chordata and Protostomia, we propose the name of Centroneuralia for this putative clade.

## Acknowledgments

We are grateful to members of Telford and Yang labs (CLOE, UCL) for discussions and comments on the manuscript. This work was funded by BBSRC grant BB/R016240/1 (PK); by a Leverhulme Trust Research Project Grant RPG-2018-302 (DJL); and by the European Union’s Horizon 2020 research and innovation programme under the Marie Skłodowska-Curie grant agreement No 764840 IGNITE (PN).

## Author Contributions

Conceptualization: MJT, PK, HP; Investigation: PK, PN, DJL, MF, NJ, RRC; Data Curation: PK, PN, DJL; Writing – Original Draft: MJT; Writing – Review & Editing: PK, PN, DJL, RRC, HP, IAR; Visualization: PK, PN, DJL, IAR, MJT; Supervision: MJT; Funding Acquisition: MJT.

## Materials and Correspondence

Correspondence and material requests should be addressed to

## Methods

### Data Availability

The simulation results, subsamples of datasets and any custom scripts used in this study are available in the GitHub repository: https://github.com/MaxTelford/MonoDeutData

### Datasets

All analyses were based on five recently published phylogenomic datasets (^2–6^) covering all major clades of the animal phylogeny. For convenience we will refer to each dataset by the name of the first author of each study (i.e., “Marletaz”, “Philippe”, “Laumer”, “Cannon’’ and “Rouse”). In all our analyses, we kept the full set of taxa apart from the fast evolving Acoelomorpha species that have been shown to be prone to phylogenetic inference errors^4^. In our analyses the Cannon dataset consisted of 67 taxa and 881 genes (“proteinortho.phy” in the original study), the Laumer dataset consisted of 422 genes and 152 taxa (167 taxa in dataset M in the original study), the Marletaz dataset consisted of 70 species and 1174 genes (alignments provided in the original study after the HMMClean and BMGE filtering), the Philippe dataset consisted of 51 taxa and 1173 genes (trimmed alignments provided in the original study) and finally the Rouse dataset consisted of 26 taxa and 1178 partitions (“70.fas” in the original study). Each dataset contained several representatives of the major metazoan clades, specifically the two Protostomia clades Lophotrochozoa and Ecdysozoa, the two Deuterostomia clades Chordates and Xenambulacraria as well as several non-bilaterian phyla.

### Measuring support for deuterostome and protostome branches

The protostomes and the deuterostomes are traditionally treated as two equally strongly supported clades. To assess whether indeed molecular phylogenetics supports this perception of bilaterian evolution we compared the branch lengths of the two clades across the five concatenated datasets as well as for each gene of each dataset individually. For each dataset we assumed the tree topology as reported in the original studies, specifically, for Cannon, Laumer, Marletaz, Rouse the trees we used relate to Fig. 2, Fig. 2a, Fig. 2, Fig. 3 from their respective publications, while for the Philippe data we assumed the relationships as in the “PHILIPPE-ALLSPP-NOACOEL-CATGTR-100JP.tre” from the original publication. For the Cannon, Laumer and Rouse data, *Xenoturbella* are considered to be sister to Ambulacraria rather than Nephrozoa. This position for *Xenoturbella* in the absence of the long branched Acoelomorpha, is supported by these datasets. For the Philippe and Marletaz data we altered the tree only to enforce deuterostome monophyly as this was not supported in the published phylogenies. For the analyses performed per gene we cropped the original trees using newick-tools^36^ such that only the taxa present in the relevant gene-alignment were present. Given the fixed topology, the branch lengths were estimated with IQ-TREE^37^ under the state frequency heterogeneous model LG+F+G model. We used the optimised topologies to measure the length of the branch leading from the common ancestor of Bilateria (protostomes plus deuterostomes) to the common ancestors respectively of deuterostomes and protostomes.

We performed a second analysis where we measured the difference in the likelihood score between the fully resolved phylogeny and for the topology after collapsing either the protostome of the deuterostome branch into a polytomy.

Collapsed Protostomia branch = (Lophotrochozoa,Ecdysozoa,(Ambulacraria,Chordata))

Collapsed Deuterostomia branch = ((Lophotrochozoa,Ecdysozoa),Ambulacraria,Chordata).

For calculating the lnLikelihoods we used the LG+F+G model. We measured the difference in lnLikelihood in both the five full concatenated alignments and for each gene from each dataset individually.

### Support for monophyletic versus paraphyletic deuterostomes and protostomes

We measured the proportion of genes in each dataset that strongly support the monophyly of the deuterostome clade “DM” over the two alternative paraphyletic topologies. Specifically, we compared the lnLikelihood of the topologies assuming monophyletic deuterotomes “DM” and two possible paraphyletic alternatives

Deuterostome Paraphyly “D1”: (Chordata,(Xenambulacraria,(Lophotrochozoa,Ecdysozoa)))
Deuterostome Paraphyly “D2”: (Xenambulacraria,(Chordata,(Lophotrochozoa,Ecdysozoa)))

We performed the equivalent analyses for the protostomes, specifically, we compared the lnLikelihood of the topologies assuming monophyletic protostomes “PM” and the two possible paraphyletic alternatives

Protostome Paraphyly “P1”: (Ecdysozoa,(Lophotrochozoa,(Xenambulacraria,Chordata)))
Protostome Paraphyly “P2”: (Lophotrochozoa,(Ecdysozoa,(Xenambulacraria,Chordata)))

To achieve this in both cases we calculated the log-likelihoods for the three alternative topologies using IQ-TREE under the LG+F+G model.

In all cases the topologies were fixed as described earlier and we manually adjusted them to produce the two alternative hypotheses rendering protostomes or deuterostomes paraphyletic. As before, each of the topologies was trimmed to the taxa present in each gene. We visualised the relative support of each gene for the three alternative topologies affecting either deuterostomes or protostomes using a method adapted from that described in ^38^. By scaling the three likelihoods in the range [0,1] such that lnLn1 + lnL2 + lnL3 = 1 (the relevant python script “likelihood_transform.py” is available in https://github.com/MaxTelford/MonoDeutData) we could use their transformed values as coordinates in a ternary plot, whose corners represent the three topologies. We considered a gene to support a particular topology if the corresponding scaled likelihood was larger than ⅔ (and the likelihoods for the other two topologies was therefore smaller than ⅓). We divided the triangle into compartments to reflect these cutoffs and plotted the points using the R package ‘ggtern’^39^ (the relevant R script “plot_triangles.R” is available in https://github.com/MaxTelford/MonoDeutData).

### Phylogenetic informativeness of genes supporting each topology

To identify potential reasons for individual genes supporting alternative topologies for both the deuterostome and the protostome clades we evaluated two parameters known to be related to errors in phylogenetic inference. For each gene alignment from each dataset we first measured the alignment length. Subsequently we inferred the Maximum Likelihood phylogeny for each gene with IQ-TREE under the LG+F+G model and measured the monophyly score of the tree (this is a measure of a data set’s ability to reconstruct known monophyletic groups and is described in ^4^). We used a Welch’s t-test to determine whether the difference in these scores between the sets of genes supporting different topologies was significant (Extended Data Table 3).

### Clade lengths across datasets

We assessed how the average branch length of the two clades Protostomes and Deuterostomes and of the outgroup clan differ across the five datasets. Using the complete alignments, we estimated the branch lengths under the LG+F+G model assuming the DM topology with IQ-TREE. To measure the average tip to stem branch length in each clade or clan we used the “pynt.py” script (available at https://github.com/MaxTelford/XenoCtenoSims). We additionally measured the tree length (sum of all branch lengths) for the five datasets after reducing each of them to the 8 species they all had in common (i.e. *Amphimedon queenslandica, Nematostella vectensis, Strongylocentrotus purpuratus, Ciona intestinalis, Peripatopsis capensis, Priapulus caudatus, Capitella teleta, Lottia gigantea*).

### Measuring branch lengths with different models and cross-validation

We compared the estimates for the deuterostome, protostome and bilaterian stem branch lengths across four models accommodating or not state composition heterogeneity across sites. We performed this analysis only for the Laumer dataset that showed the strongest support for the deuterostome monophyly. To lessen the computational burden, we reduced the dataset to 36 taxa that cover all major branches of the phylogeny selecting taxa with fewest missing data and randomly selected 50,000 of the original 106,186 alignment sites (“reduced-Laumer”). Using the reduced-Laumer dataset and assuming the DM topology we performed a phylobayes-mpi (version 1.8^10^) run for four substitution models, i.e., LG+G (homogeneous model in terms of state composition heterogeneity across sites), C10+LG+G (heterogeneous model assuming 10 different sets of amino acid state frequencies^40^), C60+LG+G (heterogeneous model assuming 60 different sets of amino acid state frequencies^40^) and the infinite mixture model CAT+LG+G model^8^. We performed 10,000 MCMC cycles for all models except for the CAT+LG+G for which we performed 20,000. For all four runs we used the final 5,000 posterior tree samples to calculate the average stem branch length of the deuterostomes, protostomes and bilaterians.

To compare how the branch length estimates relate to the fitness of each of the four models we performed cross validation analyses as described in the phylobayes-mpi manual. Using the reduced-Laumer dataset, we created 10 training datasets each consisting of 10,000 sites and 10 test datasets of 2,000 sites each. We ran phylobayes-mpi for 3,000 MCMC cycles for each of the training datasets and for each of the four models. Subsequently, using the “readpbmpi” program (distributed with the phylobayes-mpi) under the “-cv” option we calculated the posterior mean cross-validation score with a burnin of 2000 and sampling a frequency of 0.1.

### Simulations - Systematic Error

We simulated sequence alignments using parameters that match those measured from empirical sequences under the three topological hypotheses relating the deuterostome clades. As before we based these analyses on the reduced-Laumer dataset. The parameter estimates and simulations were carried out with phylobayes-mpi.

The two steps were

1. We estimated the posteriors of branch lengths and model parameters using the CAT+LG+G model on three alternative fixed topologies relating the deuterostome clades. For each fixed topology we performed 20,000 MCMC cycles with a sampling frequency of 1.
2. Using the final 5,000 posterior samples we sub-sampled with a frequency of 1 in 50, which gave us a subset of 100 posterior samples. Using these combinations of branch lengths and model parameters we simulated data with the “readpb_mpi” tool under the “ppred” option.

For phylobayes runs using all three fixed topologies, the effective sampling size (ESS), for some of the model parameters was lower than 100, but in all cases >50, possibly suggesting inadequate MCMC mixing and lack of convergence. This is a known problem of phylobayes when using large datasets. To ascertain whether the results from our simulations were affected by inadequate mixing (or by a sampling effect), we repeated the procedure for a smaller fraction (10,000 randomly selected sites) of the Laumer alignment. In this case we performed 10,000 MCMC samples, and the ESS values were above 100 for all parameters with a 25% burnin. The results of the simulations based on this dataset were in broad agreement with the results based on the larger dataset (Extended Data Table 4).

For each of the simulated datasets (simulated under the best fitting state-frequency heterogeneous CAT model) we performed two phylogenetic inference analyses using IQ-TREE, under the state-frequency homogeneous model LG+F+G and ii) the state-frequency heterogeneous model C60+LG+F+G under the PMSF approximation (Wang et al., 2018). The C60 model implements an approximation of the CAT model of Phylobayes but with 60 precomputed site frequency categories^40^ and is considerably faster to run in a maximum likelihood framework^41^, enabling the analysis of multiple simulated datasets. We wanted to test the possible influence of an LBA attracting long branched protostomes to the long branch leading to the outgroup leading to artifactual support for monophyletic deuterostomes. We repeated the previous analyses after removing the longest protostome branches (i.e., *Diuronotus aspetos*, *Geocentrophora applanata*, *Schmidtea mediterranea*, *Echinococcus multilocularis*, *Adineta vaga*, *Loa loa*, *Caenorhabditis elegans*, *Hypsibius dujardini*) and the outgroup taxa (i.e., *Polypodium hydriforme*, *Amphimedon queenslandica*, *Beroe abyssicola*, *Mnemiopsis leidyi*, *Salpingoeca rosetta*). Finally, we performed an equivalent experiment by removing the 13 shortest protostomes (*Capitella teleta*, *Helobdella robusta*, *Phyllochaetopterus* sp., *Daphnia pulex*, *Limulus polyphemus*, *Hemithiris psittacea*, *Membranipora membranacea*, *Lottia gigantea*, *Neomenia* sp., *Phoronis psammophila*, *Priapulus caudatus*) and outgroup taxa (*Craspedacusta sowerbyi*, *Nematostella vectensis*) and inferred the tree topologies under the LG+F+G model.

In all cases we measured the proportion of simulated datasets supporting each of the three potential topologies.

### Gene order

Gene order was determined from NCBI genome gff files, or gff files distributed with sequence data from the Marine Genomics Unit of Okinawa Institute of Science and Technology (https://marinegenomics.oist.jp/), or other online resources related to original publications. Orthologs of selected genes were determined from phylogenies of corresponding Pfam domains (e.g. Homeobox, Forkhead etc.). Briefly, Pfam hidden Markov models were searched against a metazoan protein database (hmmsearch), sequences were aligned (hmmalign) and then trimmed (esl-alimask, esl-alimanip) based on posterior probabilities of correct alignments and length of remaining sequence. Phylogenies were constructed using IQ-TREE, allowing the optimum model to be selected from the LG set (i.e. ‘-mset LG’ in IQ-TREE). Orthologs were identified by inspection of the phylogenetic tree around the relevant gene (e.g. *msxlx*, *pax1/9* etc.).

### Deuterostome novelties

Deuterostome novelties of Simakov et al. were searched against selected protostome and non-bilaterian metazoan data sets. The majority of gene sets (*Lingula*, *Priapulus* etc.) were obtained from the NCBI genomes resource (accessible at: https://www.ncbi.nlm.nih.gov/genome/browse#!/overview/), excepting *Phoronis australis* (OIST, see above).

Sponge transcriptomes were assembled from reads retrieved from the European Nucleotide Archive:

*P. jani*^*42*^ : SRR3417194_1.fastq.gz, SRR3417194_2.fastq.gz

*C. candelabrum*^*43*^: SRR499817_1.fastq.gz, SRR499817_2.fastq.gz, SRR499820_1.fastq.gz, SRR499820_2.fastq.gz, SRR504694_1.fastq.gz, SRR504694_2.fastq.gz

Sequences were processed into ‘left’ and ‘right’ read files adding ‘/1’ and ‘/2’ respectively to identifiers (e.g. for a ‘_1.fastq’ file: perl -pe ‘s/^@(SRR\S+)/\@$1\/1/’) and then assembled using Trinity-v2.8.4^44^ with default parameters:

trinityrnaseq-Trinity-v2.8.4/Trinity --seqType fq --max_memory 128G --left fastq/left.fastq -- right fastq/right.fastq --CPU 32

*Oscarella carmela* proteins were downloaded from compagen (http://www.compagen.org/datasets.html, file OCAR_T-PEP_130911) and *Phoronis australis* from OIST (see above).

Candidate orthologs were identified via reciprocal best hits and phylogenetic analysis of relevant Pfam domains, as described above (section ‘Gene order’).

Sequences were also searched against the NR database of the NCBI to test likely monophyly of metazoan (in this analysis generally sponge + bilaterian) proteins based on clear separation of bit scores, using blastp (v2.10.0+)^45^.

---

**Extended Data Figure 1.**
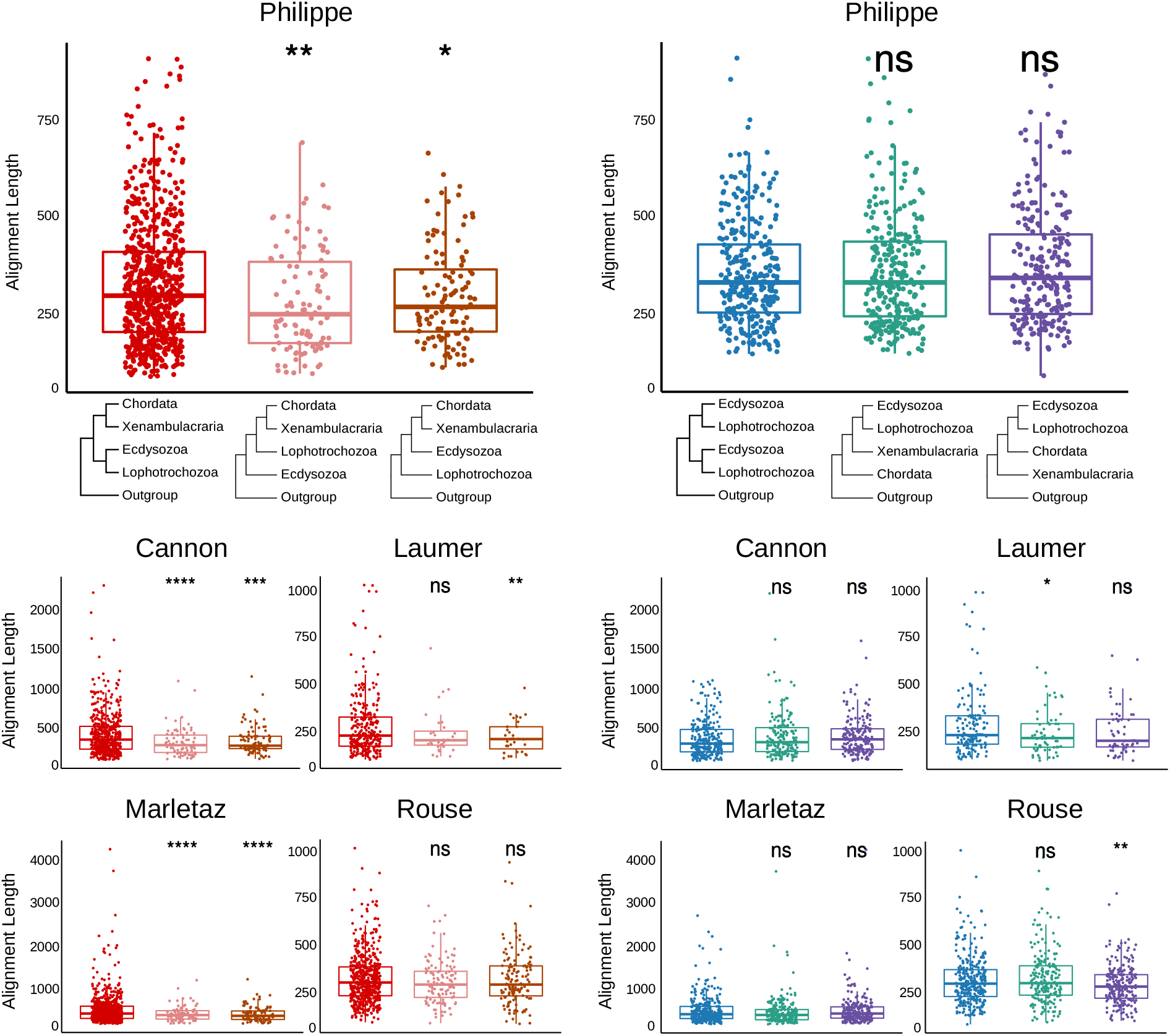
Longer genes tend to support monophyletic Protostomia but not monophyletic Deuterostomia. Box plots showing the distribution of alignment lengths for genes strongly supporting the three alternative topologies shown (sets of genes selected using the data shown in figure 2). Longer alignments are expected to contain more phylogenetic signal. Asterisks indicate significance at p < 0.05 using a Welch t-test for equal means. Left: Data from all five named datasets comparing the lengths of gene alignments supporting monophyletic Protostomia versus those supporting two alternative topologies with paraphyletic Protostomia. For most datasets the genes supporting monophyletic Protostomia are significantly longer than the genes supporting the alternative topologies. Right: Data from all five datasets comparing lengths of gene alignment supporting monophyletic Deuterostomia versus two alternative topologies with paraphyletic Deuterostomia. For most datasets the genes supporting monophyletic Deuterostomia are not significantly longer than the genes supporting the alternative topologies.

**Extended Data Table 1.** Average alignment length and monophyly scores of genes supporting the monophyly of Protostomia (PM), its alternatives (P1/P2), as well as of genes showing no preference to any of the three topologies (Px).

| Dataset | Topology | Proportion | Alignment Length | Monophyly Score |
| --- | --- | --- | --- | --- |
| Cannon | PM | 61.75% | 269.6 | 0.628 |
|  | P1 | 9.20% | 214 | 0.589 |
|  | P2 | 7.92% | 232.4 | 0.55 |
|  | Px | 14.22% | 195 | 0.568 |
| Laumer | PM | 65.88% | 408.7 | 0.564 |
|  | P1 | 7.35% | 309.3 | 0.554 |
|  | P2 | 7.58% | 310.8 | 0.559 |
|  | Px | 12.56% | 215.3 | 0.536 |
| Marletaz | PM | 61.50% | 426.2 | 0.634 |
|  | P1 | 9.20% | 346.8 | 0.594 |
|  | P2 | 7.92% | 324 | 0.584 |
|  | Px | 14.22% | 245.5 | 0.577 |
| Philippe | PM | 58.65% | 323.4 | 0.626 |
|  | P1 | 9.55% | 293.3 | 0.59 |
|  | P2 | 8.70% | 277.7 | 0.582 |
|  | Px | 14.32% | 224.9 | 0.564 |
| Rouse | PM | 46.94% | 340.3 | 0.494 |
|  | P1 | 11.46% | 339.8 | 0.474 |
|  | P2 | 10.02% | 323.8 | 0.466 |
|  | Px | 23.01% | 321.1 | 0.478 |
| **Average** | **PM** | **58.94%** | **353.6** | **0.589** |
|  | **P1** | **9.71%** | **300.6** | **0.56** |
|  | **P2** | **8.66%** | **293.7** | **0.548** |
|  | **Px** | **14.82%** | **240.4** | **0.546** |

**Extended Data Table 2.** Average alignment length and monophyly scores of genes supporting the monophyly of Deuterostomia (DM), its alternatives (D1/D2), as well as of genes showing no preference to any of the three topologies (Dx).

| Dataset | Topology | Proportion | Alignment Length | Monophyly Score |
| --- | --- | --- | --- | --- |
| Cannon | DM | 28.60% | 389 | 0.621 |
|  | D1 | 22.02% | 424.7 | 0.638 |
|  | D2 | 20.09% | 419.5 | 0.622 |
|  | Dx | 17.25% | 264.3 | 0.58 |
| Laumer | DM | 35.07% | 286.2 | 0.603 |
|  | D1 | 14.93% | 265.9 | 0.6 |
|  | D2 | 15.17% | 244.7 | 0.599 |
|  | Dx | 22.99% | 189.8 | 0.617 |
| Marletaz | DM | 27.34% | 412.3 | 0.562 |
|  | D1 | 18.99% | 407.3 | 0.58 |
|  | D2 | 20.27% | 362.6 | 0.558 |
|  | Dx | 20.7% | 240.7 | 0.533 |
| Philippe | DM | 25.58% | 309.1 | 0.617 |
|  | D1 | 20.55% | 324.8 | 0.615 |
|  | D2 | 23.02% | 314.3 | 0.612 |
|  | Dx | 21.06% | 237 | 0.577 |
| Rouse | DM | 26.66% | 350.4 | 0.48 |
|  | D1 | 29.35% | 323.6 | 0.496 |
|  | D2 | 17.32% | 362.7 | 0.496 |
|  | Dx | 24.28% | 297.1 | 0.484 |
| **Average** | **DM** | **28.65%** | **349.4** | **0.577** |
|  | **D1** | **19.17%** | **349.3** | **0.586** |
|  | **D2** | **19.17%** | **340.8** | **0.577** |
|  | **Dx** | **21.26%** | **245.8** | **0.558** |

**Extended Data Table 3.** Results of t-test comparing alignment length of genes supporting the monophyly of either Protostomia (PM) or Deuterostomia (DM) and the two alternatives (P1/P2 and D1/D2 correspondingly)

| **Dataset** | **Genes Supporting topology 1** | **Genes Supporting topology 2** | **P-value** |
| --- | --- | --- | --- |
| Laumer | **DM(148)** | **D1(64)** | 0.04945* |
| Philippe | **DM(300)** | **D1(270)** | 0.6855 |
| Cannon | **DM(252)** | **D1(177)** | 0.241 |
| Marletaz | **DM(321)** | **D1(238)** | 0.08241 |
| Rouse | **DM(314)** | **D1(204)** | 0.3317 |
| Laumer | **DM(148)** | **D2(63)** | 0.4596 |
| Philippe | **DM(300)** | **D2(241)** | 0.2372 |
| Cannon | **DM(252)** | **D2(194)** | 0.1436 |
| Marletaz | **DM(321)** | **D2(223)** | 0.8731 |
| Rouse | **DM(314)** | **D2(228)** | 0.008017* |
| Laumer | **PM(278)** | **P1(32)** | 0.1285 |
| Philippe | **PM(688)** | **P1(102)** | 0.001891* |
| Cannon | **PM(544)** | **P1(80)** | 0.00001572* |
| Marletaz | **PM(722)** | **P1(93)** | 0.00001684* |
| Rouse | **PM(553)** | **P1(118)** | 0.1573 |
| Laumer | **PM(278)** | **P2(31)** | 0.004006* |
| Philippe | **PM(688)** | **P2(112)** | 0.02411* |
| Cannon | **PM(544)** | **P2(97)** | 0.0002559* |
| Marletaz | **PM(722)** | **P2(108)** | 0.000005674* |
| Rouse | **PM(553)** | **P2(135)** | 0.9662 |

\*Statistically significantly different

**Extended Data Table 4.**
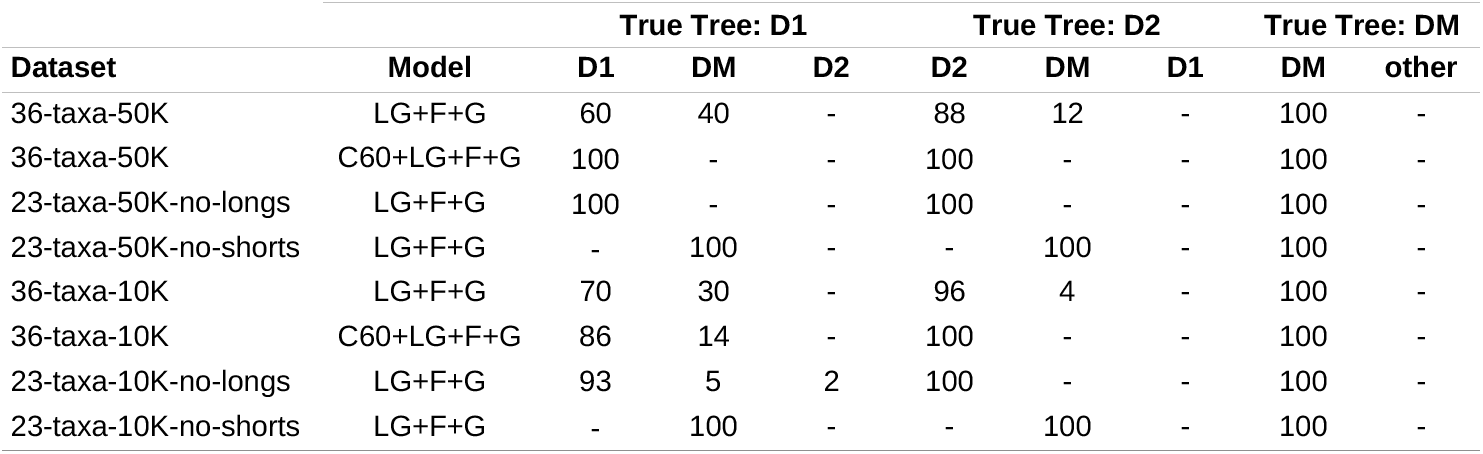
Number of simulation replicates supporting the three alternative topologies under three different true tree hypotheses (D1, D2, DM) for different datasets and models

